## Supplemental figures and tables for "The meiotic LINC complex component KASH5 is an activating adaptor for cytoplasmic dynein"

### Garner et al., Supplementary Figure 1

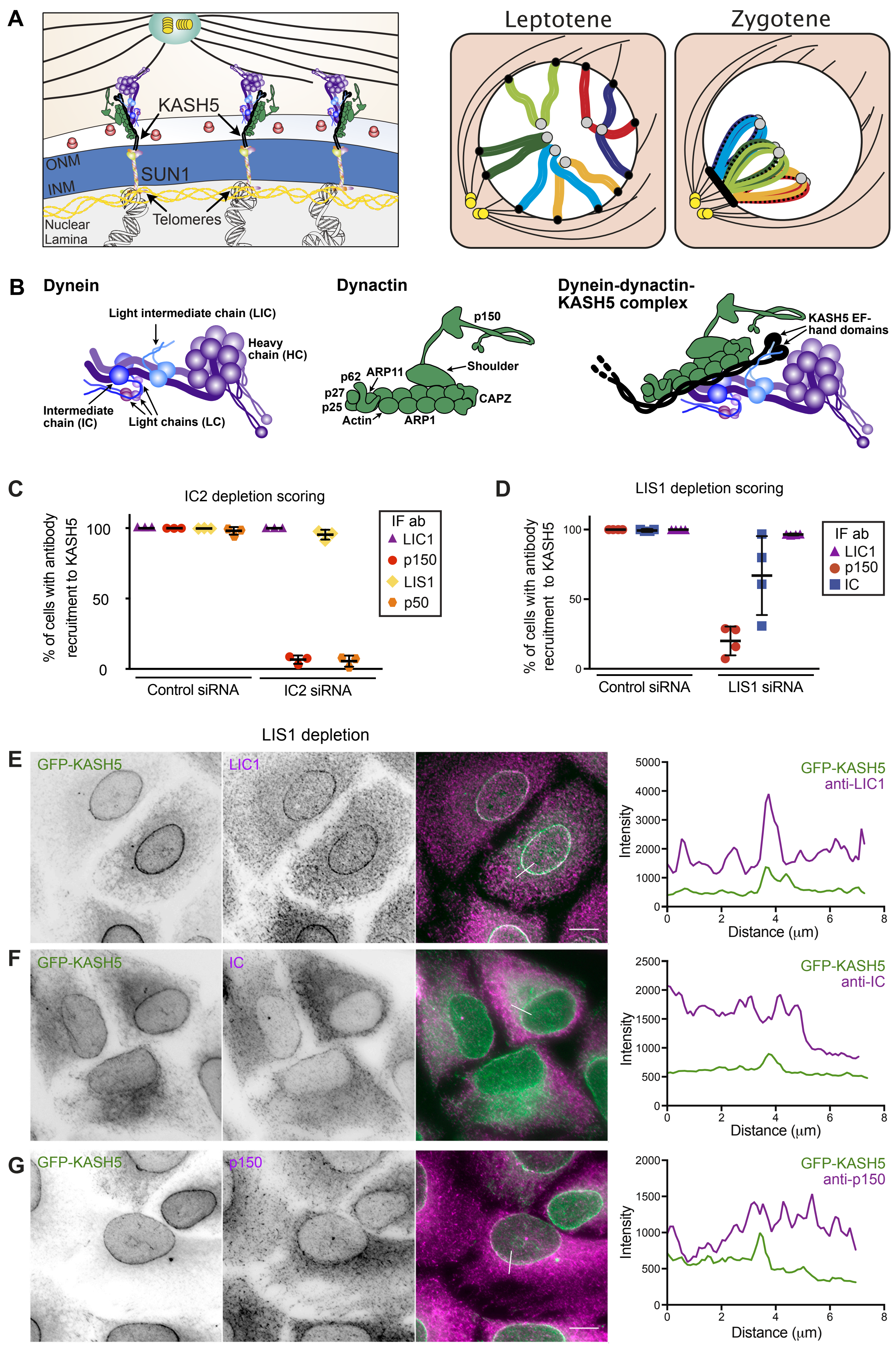

### Garner et al. Supplementary Figure 2

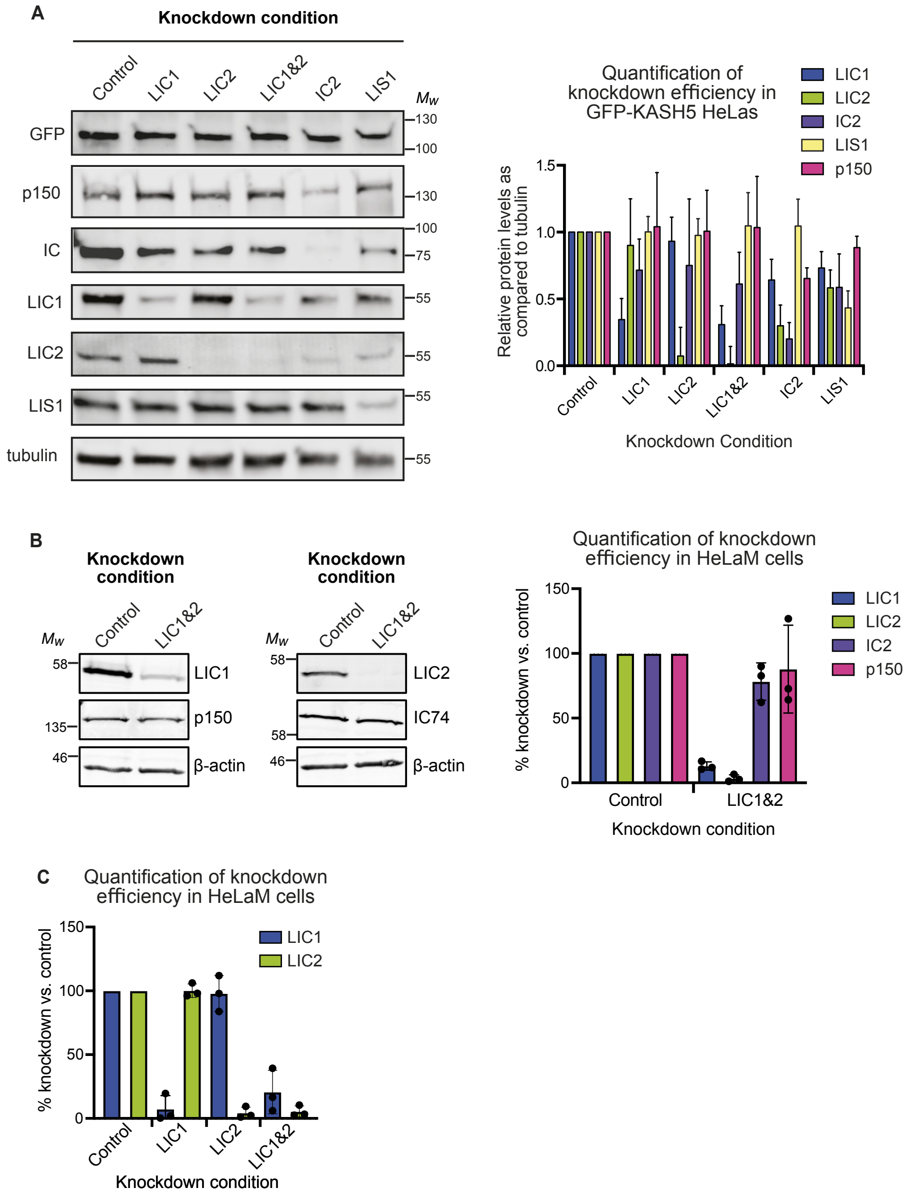

### Supplementary Figure 3, Garner et al.

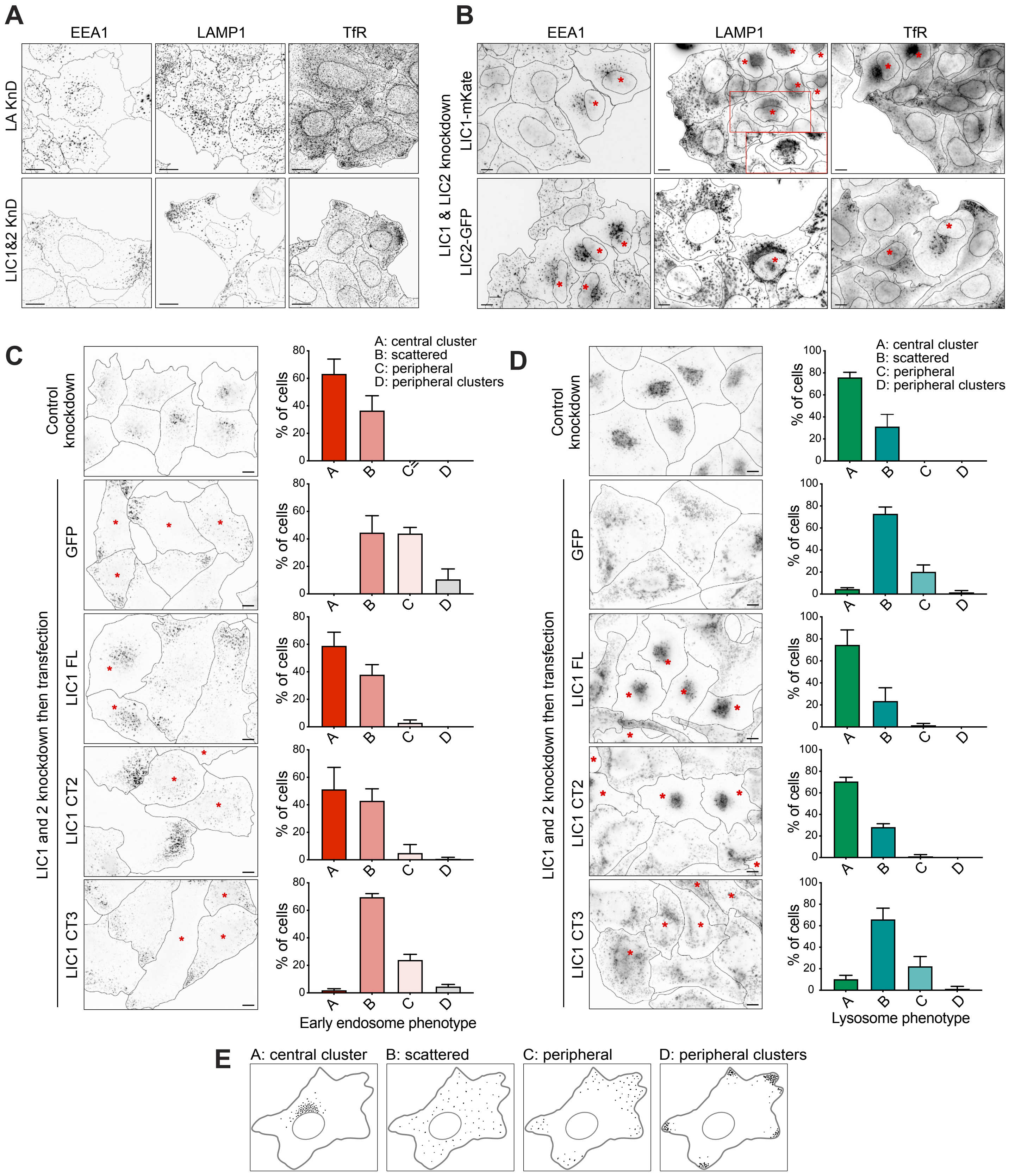

### Garner et al. Supplementary Figure 4

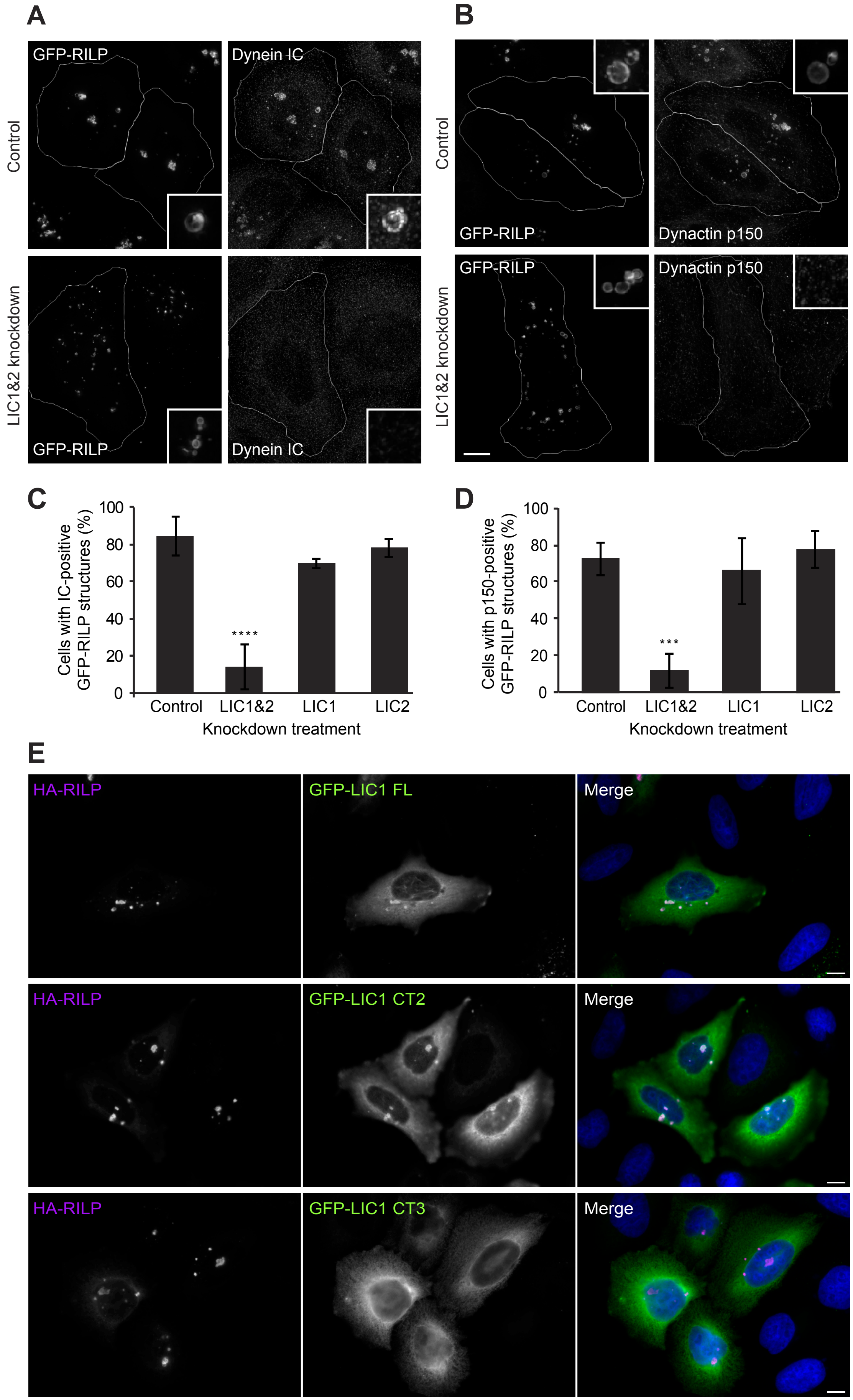

### Garner et al. Supplementary Figure 5

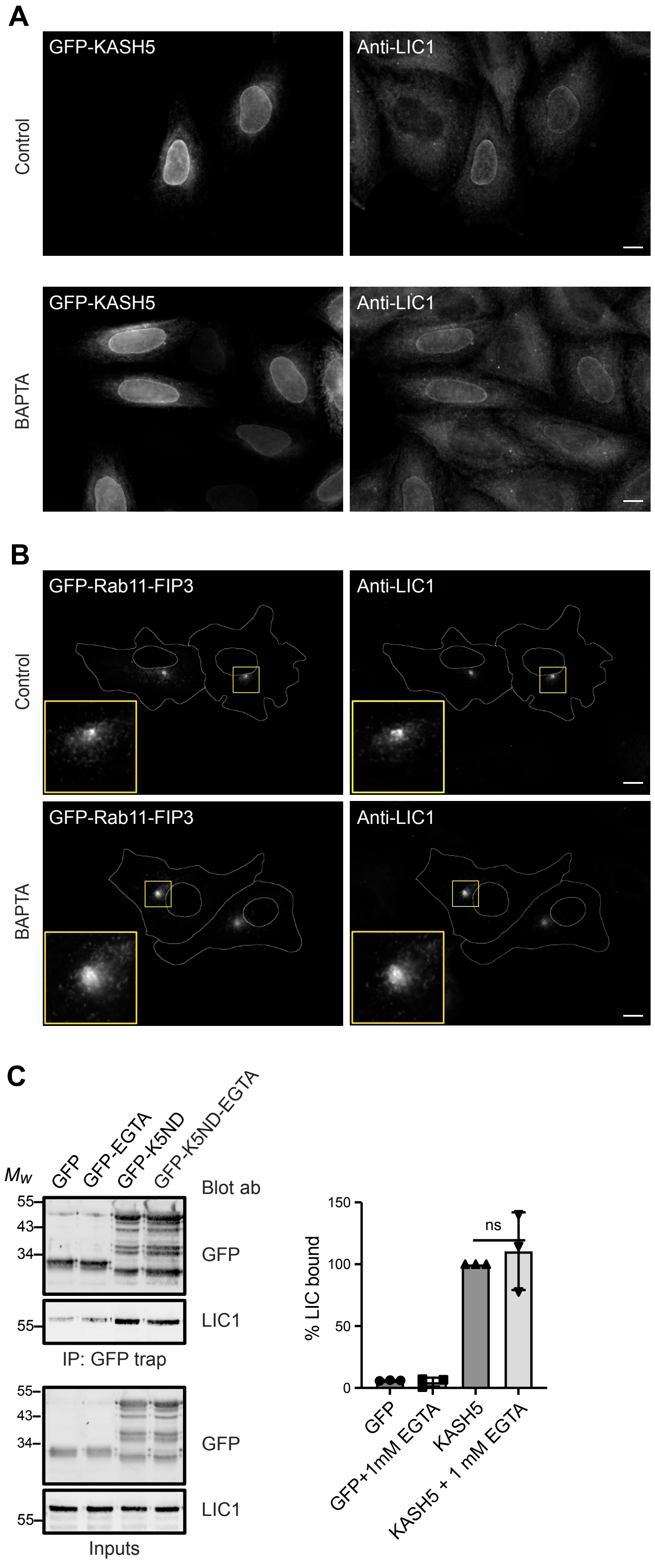

**Table S1. Quantitation of immunoblots shown in Figs 1C and 3E**

| | Fig. 1C. GFP-KASH5DK or GFP-nesprin2aDK pull-downs | | Fig. 3E. GFP-KASH5DK or GFP-nesprin2aDK pull-downs in siRNA-treated cells | | |
|---|---|---|---|---|---|
| Pull-down condition | GFP-KASH5DK | GFP-N2aDK | GFP-KASH5DK control kd | GFP-KASH5DK LIC1&2 kd | GFP-N2aDK control kd |
| IC | 103.6 | 0.1 | 100 | 3.4 | 0.2 |
| LIC1 | 74.4 | 0.4 | 100 | 1.1 | 0.1 |
| LIC2 | 78.3 | 0.6 | 100 | 0.04 | 0.1 |
| p150 | 20.4 | 0.2 | 100 | 2.9 | 2.3 |
| LIS1 | 37.1 | 1.3 | | | |
| BICD2 | 0.5 | 0.0 | | | |

For Fig. 1C, the values were calculated as (bound-prey/input-prey)/(bound-GFP/input-GFP)*100. Since the Fig. 3E data were from siRNA-treated cells the bound/input ratio of prey protein was not used. Instead, the amount of protein present in the GFP-KASH5DK pull-down in control siRNA-treated cells was set at 100%, and the prey protein present in the other pull-downs were expressed relative to that. In Fig. 3E, the knock-down efficacy for LIC1 and LIC2 was 85%.

**Table S2. Multinomial logistic regression statistical analysis of Golgi apparatus morphology following LIC depletions and rescue by GFP and GFP-LIC1 constructs.**

The data presented graphically in Figure 5C and E were analysed by multinomial logistic regression, which compares the most likely control phenotype with those observed in test samples. The odds of obtaining each phenotype, relative to the commonest normal phenotype (cluster/ribbon), was calculated for each experimental condition compared to the odds in the reference condition of either control siRNA treatment (for Figure 5C), or LIC1 and 2 double depletion followed by rescue with full length GFP-LIC1 (for Figure 5E). Analysis of 100 cells per condition, in each of 3 independent experiments. The change in odds, 95% confidence intervals and P value that the change is significant are shown in each case. Grey fill indicates non-significant comparisons.

| **Figure 5 C analysis** | **Cluster/ribbon** | **Broken ribbon** | **Semi-scatter** | **Full scatter** | **Key** |
|---|---|---|---|---|---|
| Control siRNA vs. LIC1 kd | (1.0)<br>n/a<br>n/a | 37.4<br>22-64<br><0.0001 | 5.4<br>1.7-17<br>0.004 | 3.3<br>0.7-17<br>0.144 | Change in odds<br>95% CI<br>P value |
| Control siRNA vs. LIC2 kd | (1.0)<br>n/a<br>n/a | 20.0<br>11.9-33.7<br><0.0001 | 10.4<br>3.9-28<br><0.0001 | 2.3<br>0.4-11.8<br>0.301 | Change in odds<br>95% CI<br>P value |
| Control siRNA vs. LIC 1 and 2 kd | (1.0)<br>n/a<br>n/a | 33.6<br>8-140<br><0.0001 | 601<br>137-2631<br><0.0001 | 8104<br>1617-40617<br><0.0001 | Change in odds<br>95% CI<br>P value |
| **Figure 5 E analysis** | | | | | |
| control siRNA vs. LIC kd + GFP rescue | (1.0)<br>n/a<br>n/a | 8.0<br>2.7-17<br><0.00001 | 34<br>15-76<br><0.00001 | 230<br>99-534<br><0.00001 | Change in odds<br>95% CI<br>P value |
| control siRNA vs. LIC kd + LIC1 FL rescue | (1.0)<br>n/a<br>n/a | 1.28<br>0.9-1.8<br>0.179 | 0.72<br>0.4-1.3<br>0.276 | 0.86<br>0.4-1.8<br>0.690 | Change in odds<br>95% CI<br>P value |
| control siRNA vs. LIC kd + LIC1 CT2 rescue | (1.0)<br>n/a<br>n/a | 1.05<br>0.7-1.5<br>0.800 | 1.12<br>0.7-1.9<br>0.665 | 1.24<br>0.6-2.5<br>0.540 | Change in odds<br>95% CI<br>P value |
| control siRNA vs. LIC kd + LIC1 CT3 rescue | (1.0)<br>n/a<br>n/a | 4.9<br>2.8-8.7<br><0.00001 | 10.5<br>5.5-10.1<br><0.00001 | 90<br>46-179<br><0.00001 | Change in odds<br>95% CI<br>P value |
| LIC kd with LIC1 FL rescue vs. GFP rescue | (1.0)<br>n/a<br>n/a | 6.3<br>2.9-13<br><0.00001 | 47<br>21-109<br><0.00001 | 268<br>112-640<br><0.00001 | Change in odds<br>95% CI<br>P value |
| LIC kd with LIC1 FL rescue vs. LIC1 CT2 rescue | (1.0)<br>n/a<br>n/a | 0.82<br>0.6-1.2<br>0.282 | 1.6<br>0.9-2.8<br>0.135 | 1.4<br>0.7-3.0<br>0.320 | Change in odds<br>95% CI<br>P value |
| LIC kd with LIC1 FL rescue vs. LIC1 CT3 rescue | (1.0)<br>n/a<br>n/a | 3.9<br>2.2-6.8<br><0.00001 | 14.5<br>7.3-29<br><0.00001 | 105<br>51-215<br><0.00001 | Change in odds<br>95% CI<br>P value |
| LIC kd with LIC1 CT2 rescue vs. control siRNA | (1.0)<br>n/a<br>n/a | 0.78<br>0.5-1.1<br>0.179 | 1.4<br>0.8-2.5<br>0.276 | 1.2<br>0.6-2.4<br>0.690 | Change in odds<br>95% CI<br>P value |

**Table S3. Multinomial logistic regression statistical analysis of Golgi apparatus morphology following expression of GFP-KASH5ΔK wild-type and EF-hand mutants.**

The data presented graphically in Figure 8E were analysed by multinomial logistic regression, which compares the most likely control phenotype with those observed in test samples. The odds of obtaining each phenotype, relative to the commonest normal phenotype, was calculated for each experimental condition compared to the odds in the reference condition. The reference conditions were the commonest normal Golgi apparatus phenotype (cluster/ribbon), in either GFP or GFP-KASH5ΔK-AA expressing cells. Analysis of 100 cells per condition, in each of 3 independent experiments. The change in odds, 95% confidence intervals and P value that the change is significant are shown in each case. Grey fill indicates non-significant comparisons.

| Condition | Cluster/ribbon | Broken ribbon | Semi-scatter | Full scatter | Key |
|---|---|---|---|---|---|
| GFP vs.<br>GFP-KASH5ΔK-WT | (1.0)<br>n/a<br>n/a | 17.2<br>4.4-66.5<br><0.0001 | 362<br>95-1388<br><0.0001 | 3796<br>684-21073<br><0.0001 | Odds ratio<br>95% CI for odds<br>P value |
| GFP vs.<br>GFP-KASH5ΔK-*fue* | (1.0)<br>n/a<br>n/a | 5.6<br>3.8-8.3<br><0.0001 | 6.1<br>3.5-10.5<br><0.0001 | 7.4<br>1.9-29<br>0.004 | Odds ratio<br>95% CI for odds<br>P value |
| GFP vs.<br>GFP-KASH5ΔK-AA | (1.0)<br>n/a<br>n/a | 5.1<br>3.4-7.5<br><0.0001 | 7.7<br>4.5-13.2<br><0.0001 | 7.4<br>1.8-29.2<br>0.004 | Odds ratio<br>95% CI for odds<br>P value |
| GFP vs.<br>GFP-KASH5ΔK-Mod 1 | (1.0)<br>n/a<br>n/a | 4.8<br>3.3-7.1<br><0.0001 | 6.1<br>3.6-10.6<br><0.0001 | 4.6<br>1.1-19.9<br>0.040 | Odds ratio<br>95% CI for odds<br>P value |
| GFP vs.<br>GFP-KASH5ΔK-Mod 2 | (1.0)<br>n/a<br>n/a | 6.6<br>1.8-24.3<br>0.004 | 201<br>59.2-683<br><0.0001 | 4142<br>823-20859<br><0.0001 | Odds ratio<br>95% CI for odds<br>P value |
| GFP-KASH5ΔK-AA vs.<br>GFP-KASH5ΔK-*fue* | (1.0)<br>n/a<br>n/a | 1.1<br>0.7-1.7<br>0.643 | 0.8<br>0.5-1.3<br>0.366 | 1.0<br>0.3-3.0<br>0.995 | Odds ratio<br>95% CI for odds<br>P value |
| GFP-KASH5ΔK-AA vs.<br>GFP-KASH5ΔK-Mod 1 | (1.0)<br>n/a<br>n/a | 1.3<br>0.4-4.8<br>0.694 | 26.1<br>7.8-87<br><0.0001 | 562<br>142-2232<br><0.0001 | Odds ratio<br>95% CI for odds<br>P value |
| GFP-KASH5ΔK-AA vs.<br>GFP-KASH5ΔK-Mod 2 | (1.0)<br>n/a<br>n/a | 1.0<br>0.6-1.4<br>0.832 | 0.8<br>0.5-1.3<br>0.381 | 0.6<br>0.2-2.1<br>0.443 | Odds ratio<br>95% CI for odds<br>P value |
| GFP-KASH5ΔK-AA vs.<br>GFP | (1.0)<br>n/a<br>n/a | 0.2<br>0.1-0.3<br><0.0001 | 0.13<br>0.08-0.2<br><0.0001 | 0.14<br>0.03-0.54<br>0.004 | Odds ratio<br>95% CI for odds<br>P value |
| GFP-KASH5ΔK-AA vs.<br>GFP-KASH5ΔK-WT | (1.0)<br>n/a<br>n/a | 3.4<br>0.9-13.2<br>0.081 | 47<br>12-177<br><0.0001 | 515<br>116-2291<br><0.0001 | Odds ratio<br>95% CI for odds<br>P value |

**Table S4. Multinomial logistic regression statistical analysis of changes in early endosome and lysosome distribution after LIC depletion and rescue by GFP and GFP-LIC1 constructs.**

The data presented graphically in Figure S3 C and D were analysed by multinomial logistic regression, which compares the most likely control phenotype with those observed in test samples. The odds of obtaining each phenotype, relative to the commonest normal phenotype, was calculated for each experimental condition compared to the odds in the reference condition. The reference conditions were the commonest normal phenotype (clustered) in either control siRNA treatment, or LIC1 and 2 double depletions followed by rescue with full length GFP-LIC1. Analysis of 100 cells per condition, in each of 3 independent experiments. The change in odds, 95% confidence intervals and P value that the change is significant are shown in each case. Grey fill indicates non-significant comparisons.

| Figures S3C | Central endosome cluster | Scattered endosomes | Peripheral endosomes | Peripheral endosome cluster | Key |
|---|---|---|---|---|---|
| Control siRNA vs. LIC kd with GFP rescue | (1.0)<br>n/a<br>n/a | 64.8<br>20-210<br><0.0001 | 4851<br>950-14759<br><0.0001 | 1410<br>269-7394<br><0.0001 | Change in odds<br>95% CI<br>P value |
| Control siRNA vs. LIC kd with FL LIC1 rescue | (1.0)<br>n/a<br>n/a | 0.56<br>0.4-0.8<br>0.001 | 0.82<br>0.2-4.1<br>0.812 | 0.82<br>0.16-4.1<br>0.809 | Change in odds<br>95% CI<br>P value |
| Control siRNA vs. LIC kd with LIC1 CT2 rescue | (1.0)<br>n/a<br>n/a | 1.47<br>1.0-2.0<br>0.001 | 6.45<br>1.8-22.8<br>0.004 | 1.7<br>0.4-7.9<br>0.478 | Change in odds<br>95% CI<br>P value |
| Control siRNA vs. LIC kd with LIC1 CT3 rescue | (1.0)<br>n/a<br>n/a | 64.5<br>28-150<br><0.0001 | 956<br>230-3970<br><0.0001 | 195<br>43-874<br><0.0001 | Change in odds<br>95% CI<br>P value |
| LIC kd with FL LIC1 rescue vs. LIC1 CT2 rescue | (1.0)<br>n/a<br>n/a | 2.6<br>18-3.7<br><0.0001 | 7.8<br>2.2-28<br>0.001 | 2.1<br>0.46-9.7<br>0.333 | Change in odds<br>95% CI<br>P value |
| LIC kd with FL LIC1 rescue vs. LIC1 CT3 rescue | (1.0)<br>n/a<br>n/a | 115<br>49-270<br><0.0001 | 1163<br>280-4829<br><0.0001 | 238<br>53-1068<br><0.0001 | Change in odds<br>95% CI<br>P value |
| LIC kd with FL LIC1 rescue vs. GFP rescue | (1.0)<br>n/a<br>n/a | 115<br>35-376<br><0.0001 | 5899<br>1155-30129<br><0.0001 | 1721<br>328-9036<br><0.0001 | Change in odds<br>95% CI<br>P value |
| LIC kd with FL LIC1 rescue vs. control siRNA | (1.0)<br>n/a<br>n/a | 1.8<br>1.2-2.5<br>0.001 | 1.2<br>0.24-6.1<br>0.812 | 1.2<br>0.24-6.2<br>0.809 | Change in odds<br>95% CI<br>P value |
| **Figure S3E** | **Central lysosome cluster** | **Scattered lysosomes** | **Peripheral lysosomes** | **Peripheral lysosome cluster** | **Key** |
| Control siRNA vs. LIC kd with GFP rescue | (1.0)<br>n/a<br>n/a | 38<br>21-68<br><0.0001 | 331<br>92-1190<br><0.0001 | 32<br>7.3-144<br><0.0001 | Change in odds<br>95% CI<br>P value |
| Control siRNA vs. LIC kd with FL LIC1 rescue | (1.0)<br>n/a<br>n/a | 0.77<br>0.5-1.1<br>0.149 | 2.04<br>0.5-8.3<br>0.318 | 1.0<br>0.2-5.1<br>0.985 | Change in odds<br>95% CI<br>P value |
| Control siRNA vs. LIC kd with LIC1 CT2 rescue | (1.0)<br>n/a<br>n/a | 0.97<br>0.69-1.4<br>0.870 | 1.8<br>0.4-7.6<br>0.427 | 1.1<br>0.2-5.4<br>0.931 | Change in odds<br>95% CI<br>P value |

| | | | | | |
|---|---|---|---|---|---|
| Control siRNA vs. LIC kd with LIC1 CT3 rescue | (1.0)<br>n/a<br>n/a | 15.5<br>9.9-14<br><0.0001 | 165<br>49-555<br><0.0001 | 14.7<br>3.5-62<br><0.0001 | Change in odds<br>95% CI<br>P value |
| LIC kd with FL LIC1 rescue vs. LIC1 CT2 rescue | (1.0)<br>n/a<br>n/a | 1.3<br>0.9-1.8<br>0.209 | 0.9<br>0.27-2.9<br>0.835 | 1.1<br>0.2-5.3<br>0.947 | Change in odds<br>95% CI<br>P value |
| LIC kd with FL LIC1 rescue vs. LIC1 CT3 rescue | (1.0)<br>n/a<br>n/a | 20.2<br>12.7-32<br><0.0001 | 81<br>32-202<br><0.0001 | 14.5<br>3.4-61<br><0.0001 | Change in odds<br>95% CI<br>P value |
| LIC kd with FL LIC1 rescue vs. GFP rescue | (1.0)<br>n/a<br>n/a | 49<br>27-90<br><0.0001 | 162<br>60-440<br><0.0001 | 32<br>7.2-141<br><0.0001 | Change in odds<br>95% CI<br>P value |
| LIC kd with FL LIC1 rescue vs. control siRNA | (1.0)<br>n/a<br>n/a | 1.3<br>0.9-1.9<br>0.149 | 0.5<br>0.1-2.0<br>0.318 | 1.0<br>0.2-4.9<br>0.985 | Change in odds<br>95% CI<br>P value |
